# An Effective Surface Passivation Assay for Single-Molecule Studies of Chromatin and Topoisomerase II

**DOI:** 10.1101/2024.09.25.614989

**Authors:** Tung T. Le, Xiang Gao, Seong Ha Park, Jaeyoon Lee, James T. Inman, Michelle D. Wang

## Abstract

For single-molecule studies requiring surface anchoring of biomolecules, a poorly passivated surface can result in alterations of biomolecule structure and function that can result in artifacts. This protocol describes surface passivation and sample chamber preparation for mechanical manipulation of chromatin fibers and characterization of topoisomerase II activity in physiological buffer conditions. The method employs enhanced surface hydrophobicity and purified blocking proteins to reduce non-specific surface adsorption. This method is accessible, cost-effective, and potentially widely applicable to other biomolecules.

For a complete list of publications that employ this protocol, see the paper references.

**GRAPHICAL ABSTRACT:** 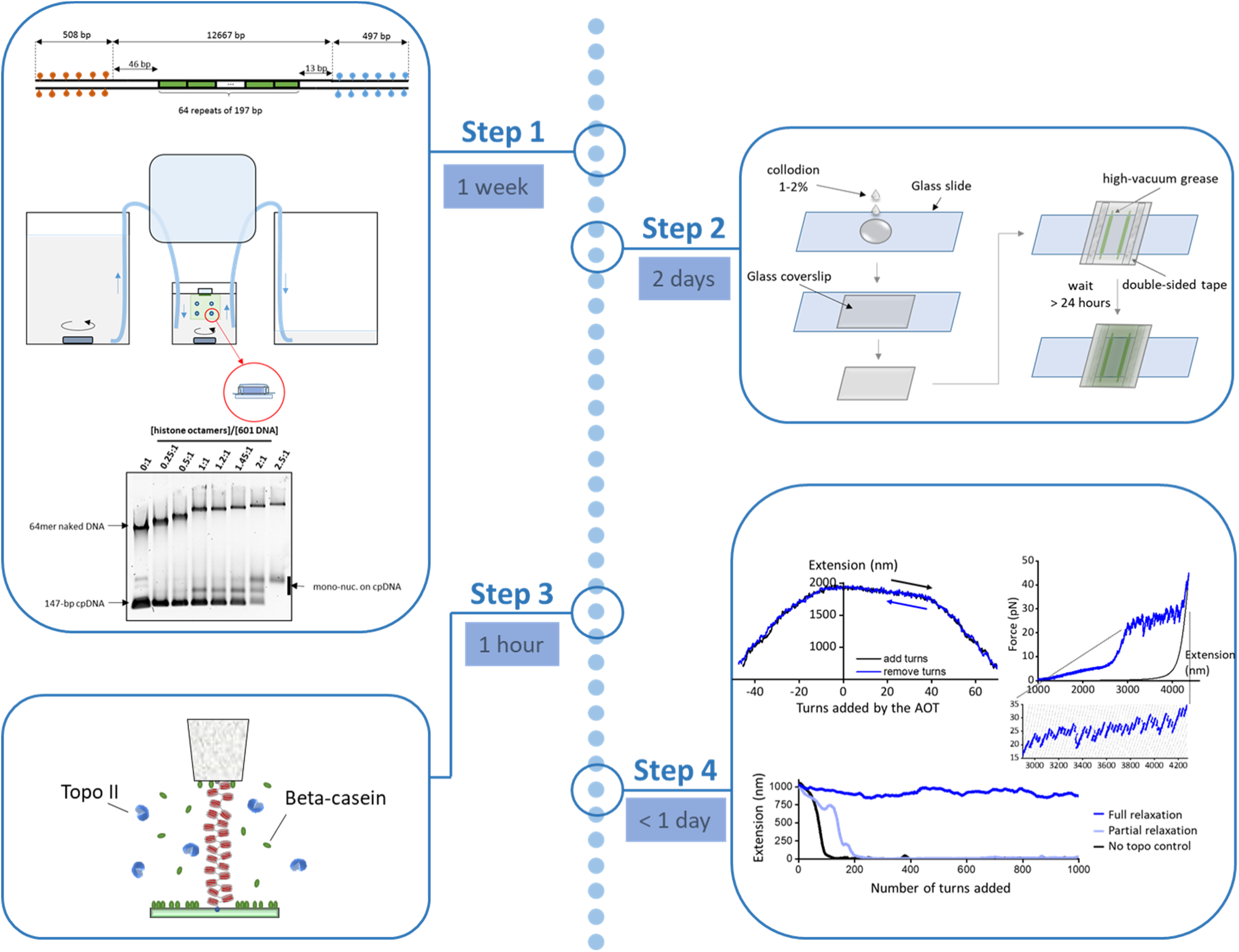

## C. BEFORE YOU BEGIN

Single-molecule experiments with biological macromolecules often require maintenance of the biomolecule’s native conformations and minimization of the interference from the confining environment. For a surface-anchored molecule, it is desirable to have minimum non-specific surface adsorption as this may increase background signal (for fluorescence imaging), induce conformational changes of the molecules, or render the protein concentration in solution uncertain, preventing the acquisition of meaningful data. For large protein complexes such as chromatin, non-specific surface adsorption was a significant bottleneck in early single-molecule studies^2^, especially when using highly saturated nucleosome arrays with minimum naked DNA in physiological buffer conditions containing Mg^2+^ at millimolar-scale concentrations. Previously, common surface-blocking reagents have been employed for chromatin studies, including mixed milk protein^3–5^, BSA^6–8^ and PEG^9–11^. However, the size of the nucleosome arrays studied was limited due to insufficient surface passivation. We developed an effective surface passivation protocol using beta-casein that enables studying long nucleosome arrays in a physiological buffer condition with minimal non-specific surface adsorption^1^. This method also proved to be effective in studying other proteins that act on DNA/chromatin substrates, such as topoisomerase II^1,12,13^.

In this protocol, we provide detailed instructions for preparing a DNA template, assembling nucleosome arrays, validating the nucleosome array quality, preparing nitrocellulose chambers, generating fiducial markers, preparing a beta-casein solution, and passivating the surface with beta-casein.

### Torsionally-constrained (TC) 64-mer DNA preparation

#### Timing: 4 days

The torsionally constrained (TC) 64-mer DNA template is formed by ligating two 500-bp multi-labeled adapters, one to each end, to a DNA template consisting of 64 tandem repeats of a 197 bp sequence, each repeat containing a 601-nucleosome positioning element (NPE)^14^ (Figure 1A). The 64-mer DNA is constructed by using a method similar to that previously described^15^. The 500-bp adapter sequence is constructed by concatenating three 145-bp sequences that were found to have low affinity for nucleosomes^16^ to prevent additional nucleosomes from assembling on the adapters. The low nucleosome affinity sequence is flanked by a set of restriction sequences so that it can be employed in various constructs.

1. Obtain the 64-mer DNA from P197NRL-64ex:

a. To generate the P197NRL-64ex plasmid (a.k.a. pMDW108), transform the plasmid into NEB *E. coli* stable competent cells according to the manufacturer’s protocol. Plate the cells, grow colonies, select individual colonies in liquid growth media, and grow cultures overnight. Purify the plasmids using a Qiagen Qiaprep kit following the manufacturer’s instructions. Check the concentration of the plasmid using a spectrophotometer.
*CRITICAL*: Care must be taken to minimize a change in NPE repeats. Keep the plasmid sample from each independent colony separate for size screening. The plasmid can lose or gain repeats during transformation, and thus, it is essential to check its size using the following procedure. For this reason, it is also advisable to store the plasmid in the form of purified DNA and not in the form of transformed cells.

b. Digest each plasmid sample with BstXI and BglI and run the resulting digestion on a 0.8% agarose gel. Run the gel at ∼100 V for 2 hours to select the plasmid with the correct size (12,667 bp) and highest homogeneity (green arrows, Figure 1B). Do not select the samples with slightly shorter fragments (red arrows, Figure 1B), indicative of the loss of the 601 repeats.
c. Pool the high-quality plasmid samples and double digest with BstXI and BglI. The protocol for the p197NRL-64ex digestion at large scale (230 µL, ∼ 71 ug plasmid) is listed below:

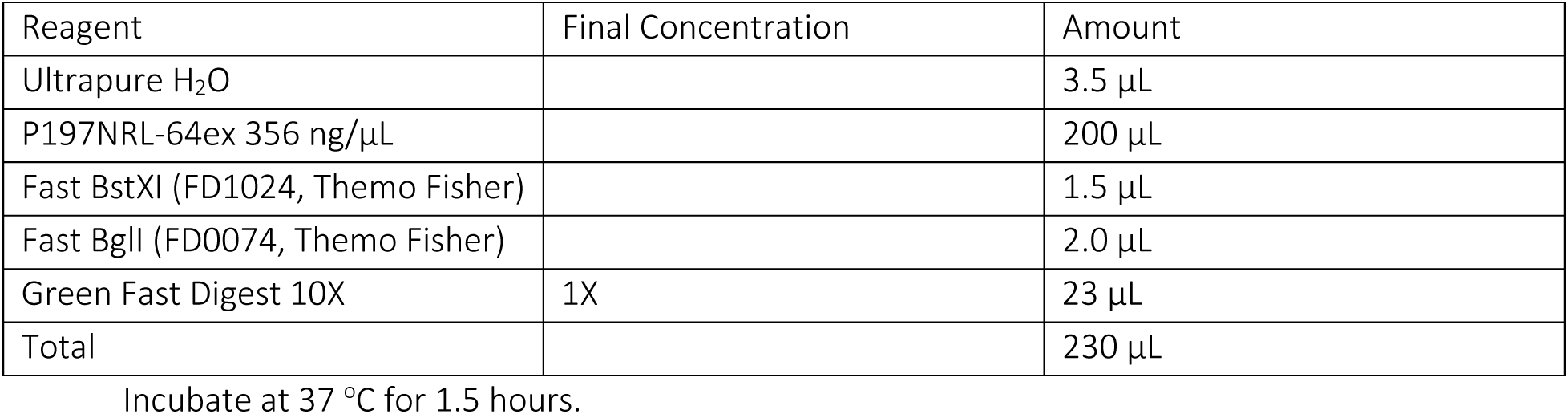

*Note:* After 45 min of enzymatic digestion, quickly check the DNA (∼ 100 ng) on a 0.8% agarose gel to confirm the digestion. If the digestion is incomplete, proceed with the full-time course. Confirm that the top band, which corresponds to the 12,667 bp 64-mer DNA, looks sharp and the other smaller DNA fragments, which are 730 bp, 570, and 558 bp, are of correct size (Figure 1C).

d. After digestion is confirmed, directly load the entire digested DNA sample into a single, large well (that can hold 300-400 µL) in a 10-cm 0.8% agarose gel and run for 40 min at 90V to distinguish the bands. For longer gels, increase the applied voltage to maintain ∼ 9V/cm.

*Note:* One can use heat-resistant tape to cover and combine multiple well combs to make a larger well.

e. Carefully excise the top 12.7 kb DNA band that contains the 64-mer DNA without shining UV light on the DNA. Excise a minimal gel size that still contains all DNA bands to facilitate DNA elution (Figure 1C). Elute the DNA using Zymoclean Large Fragment Recovery kit, following the manufacturer’s instructions.

2. DNA adapter end digestion:

a. Prepare the 500-bp DNA adapters labeled with 25% biotin tag or 25% digoxygenin tag as described in the section “Materials and Equipment” (Figure 1D).
b. Mix the following ingredients in separate 1.5-mL tubes. Incubate at 37 °C for 2.5 hours.

Digest biotin-labeled adapter with BglI

| Reagent | Final Concentration | Amount |
| --- | --- | --- |
| Ultrapure H<sub>2</sub>O | | 6.0 μL |
| Purified DNA adapter 5.8 μg | | 99.0 μL |
| Fast Digest 10X | 1X | 12.0 μL |
| Fast BglI (FD0074, Thermo Fisher) |  | 5.5 µL |
| Total |  | 120 µL |

Digest dig-labeled adapter with BstXI

| Reagent | Final Concentration | Amount |
| --- | --- | --- |
| Ultrapure H<sub>2</sub>O |  | 3.5 µL |
| Purified DNA adapter 3.1 ug |  | 99.0 µL |
| Fast Digest 10X | 1X | 12.0 µL |
| Fast BstXI (FD1024, Thermo Fisher) |  | 3.0 µL |
| Total |  | 120 µL |

b. Perform a spin column purification using PureLink PCR columns with an elution buffer (10 mM Tris-Cl pH 8.0 and 0.1 mM EDTA) prewarmed to 50 °C. Use a NanoDrop to measure DNA concentration after spin column purification.

*Note:* Adjust the elution buffer volume to obtain similar DNA concentrations of the two DNA adapters.

3. Ligate the purified 64-mer DNA with the two 500-bp sticky adapters with T4 DNA ligase.

a. Use a molar ratio: [64-mer DNA]:[biotin adapter]:[dig adapter] = 1.0 : 1.25 : 1.25

*CRITICAL:* Do not use a high molar ratio of DNA adapters vs. the 64-mer DNA (>1.5:1) in the ligation mixture as ligation between the two adapters can happen. This will result in very short DNA that can with the 64-mer DNA on the surface and bead tethering.

| Reagent | Final Concentration | Amount |
| --- | --- | --- |
| Ultrapure H<sub>2</sub>O |  | 17.5 µL |
| Purified 64-mer 341 ng/µL, 40.8 nM | 19.5 nM | 57 µL |
| 500-bp biotin-labeled adapter 190 nM | 24.5 nM | 15.5 µL |
| 500-bp dig-labeled adapter 190 nM | 24.5 nM | 15.5 µL |
| T4 10X buffer | 1X | 12 µL |
| T4 ligase 400 units/µL | ~ 8 units/µL | 2.5 µL |
| Total |  | 120 µL |

b. Keep the tube on ice and mix carefully using a wide bore tip to avoid introducing nicks to the 64-mer DNA. Incubate the samples at 16 °C for 8-10 hours, then heat-kill the ligase for 30 min at 65 °C.

*Note:* A preliminary small-scale ligation reaction (20 µL with ∼ 2-5 nM DNA) before proceeding with the large-scale ligation may help avoid wasting reagents in case there is an issue with the adapters or the 64-mer DNA. If the small-scale ligation fails, it is most likely due to poor DNA end cleavage efficiency.

4. After the ligation, immediately perform a spin-column purification using Pure Link PCR columns. Elute the DNA using an elution buffer containing 10 mM Tris-Cl pH 8.0 and 1 mM EDTA and store at 4 °C.

**Figure 1:**
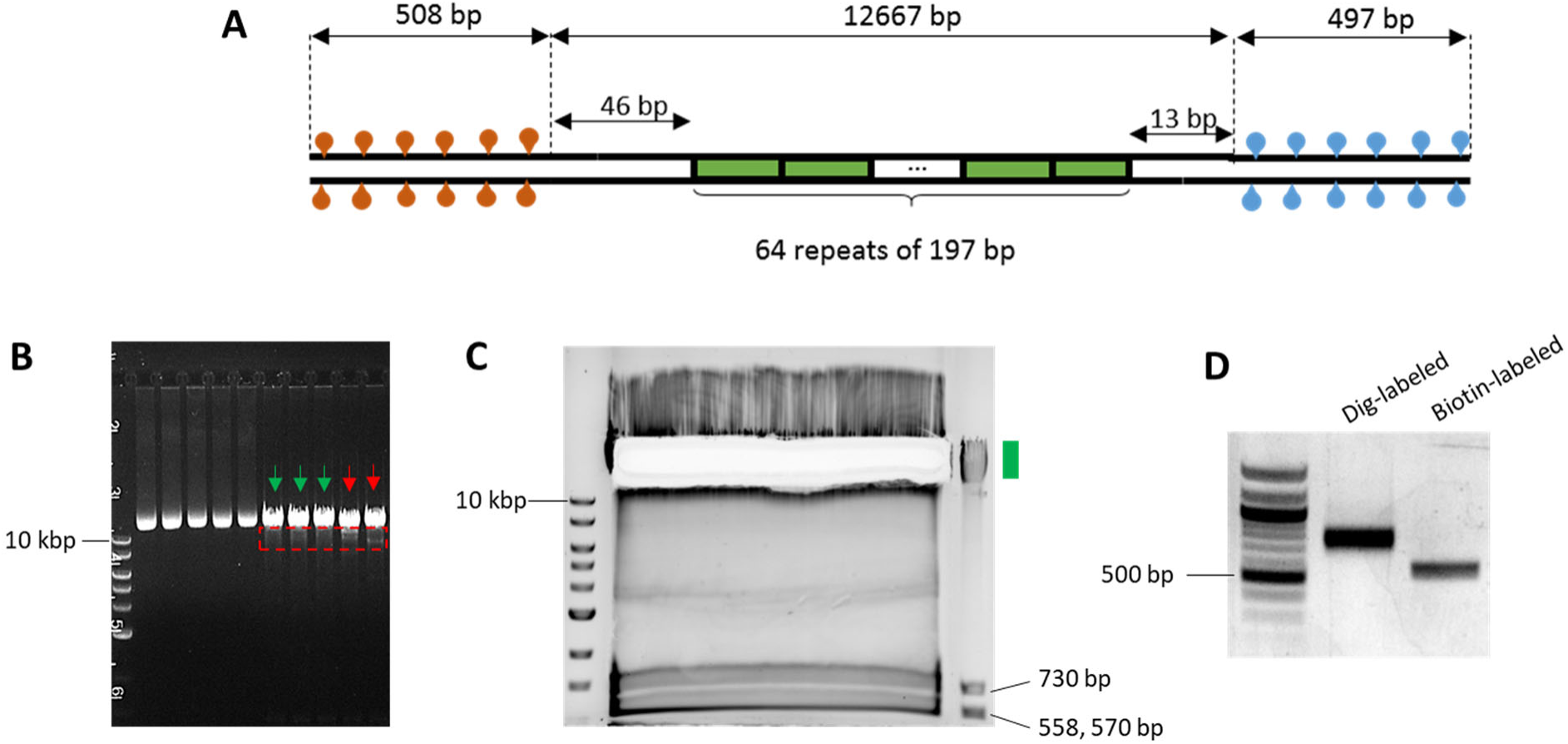
(A) DNA construct of the multiple-labeled 64-mer 601 DNA template. The two ends of the 64-mer DNA template are labeled with either biotin or digoxygenin (∼ 500 bp each), enabling the formation of a torsionally constrained DNA tether anchored between an anti-digoxygenin-coated surface and streptavidin-coated bead. (B) Selection of p197NRL-64ex plasmid with minimal 601 repeat loss (green arrows). Red arrows indicate samples with more 601 repeat loss (dashed red box). (C) Gel purification of the 64-mer DNA. The green box denotes the gel location of the desired sample. (D) Gel electrophoresis of the digoxygenin-tagged and biotin-tagged 500-bp adapters

### Nucleosome assembly

#### Timing: 2 days

In brief, nucleosome arrays are assembled onto the 64-mer DNA construct by gradient NaCl dialysis^4,17^ from 2.0 M to 0.6 M over 18 hours at 4 °C at different molar ratios (0.25:1 to 2.5:1) of histone octamers to 601 DNA repeats. An equal mass of 147-bp non-601-sequence competitor DNA to the 64-mer DNA construct is added to the reconstitution reactions to avoid nucleosome over-assembly^18^.

1. Use a razor blade to cut multiple thermal-resistant 200-µL PCR tubes into separate tube caps and tube rims (Figure 2A).
2. Collect, clean, and autoclave glassware:

a. 2 x 500-mL glass beakers, 1 x 2000-mL glass beaker, 1 x 100-mL plastic graduated cylinder
b. Rinse the glassware with ultrapure H_2_0, gently clean with diluted soap, and rinse with ultrapure H_2_0 to remove soap residues
c. Autoclave the glassware, the PCR-tube caps, and the PCR-tube rims using the ‘Fast’ cycle for 30 minutes.
3. Prepare the High-salt, Low-salt, and Zero-salt dialysis buffers as described in the section “Materials and Equipment”.
4. Move the beakers with High-salt, Low-salt, and Zero-salt dialysis buffers into the 4 °C. Add magnetic stir bars to each beaker of buffer to completely mix the solutions.
5. Assemble the incoming and outgoing tubings that are connected to the dialysis pump into the Low-salt and High-salt cylinders, respectively.
*Note:* Flush the tubing with diluted NaN_3_ (see “Materials and Equipment”) before assembly to reduce the chance of bacterial contamination.
6. Cut a piece of the dialysis membrane (MWCO: 6-8 kD) and submerge it in a Petri dish with ultrapure H_2_O for at least 10 minutes. One piece of dialysis membrane can accommodate several different assemblies.
7. Cut the edge of the dialysis membrane using a clean pair of scissors and use a pair of tweezers to carefully open the membrane without tearing it.
8. Take an aliquot of histone octamers from the -80 °C and thaw at 4 °C or on ice. After there are no crystals left, gently pipette the solution up and down to mix the histone octamers.
9. Mix the 64-mer DNA ligated with biotin and dig adapters prepared in the previous section with histone octamers following this protocol. Use a 500-µL tube and keep the tube on ice during the process. See “Materials and Equipment” for the preparation of the 2X nucleosome assembly buffer and the 147-bp DNA competitor.

| [Histone Octamers]/[601 DNA] | 1:1 | 0.5:1 | 0.25:1 |
| --- | --- | --- | --- |
| Ultrapure $\text{H}_2\text{O}$ | 3.34 $\mu\text{L}$ | 4.68 $\mu\text{L}$ | 5.36 $\mu\text{L}$ |
| Nucleosome assembly buffer 2X | 15 $\mu\text{L}$ | 15 $\mu\text{L}$ | 15 $\mu\text{L}$ |
| 64-mer DNA 213 ng/$\mu\text{L}$ (22.73 nM) | 5 $\mu\text{L}$ | 5 $\mu\text{L}$ | 5 $\mu\text{L}$ |
| 147-bp DNA competitor 268 ng/$\mu\text{L}$ | 3.97 $\mu\text{L}$ | 3.97 $\mu\text{L}$ | 3.97 $\mu\text{L}$ |
| Histone octamers 2.7 $\mu\text{M}$ | 2.69 $\mu\text{L}$ | 1.35 $\mu\text{L}$ | 0.67 $\mu\text{L}$ |
| **Total** | **30 $\mu\text{L}$** | **30 $\mu\text{L}$** | **30 $\mu\text{L}$** |

10. Discard the H_2_0 in the Petri dish containing the dialysis membrane. Drop 500 µL of the High-salt buffer onto the membrane to completely soak the membrane. Discard any excess buffer outside the membrane by carefully tilting the dish and pipetting it off to prevent excess buffer from diluting the dialysis reaction. Close the lid of the Petri dish to avoid drying the membrane.
11. Transfer ∼ 30 µL of each histone-DNA molar ratio mixture into a separate PCR-tube cap. Place the dialysis membrane on top of the filled cap. Place a PCR-tube rim over the membrane on top of each filled cap and firmly press down using a clean 1.5-mL PCR tube to form dialysis chambers (Figure 2A). Repeat the process for each assembly sample attached to the same membrane. Ensure caps are placed at least several centimeters apart to facilitate the cap assembling process. Move to a new membrane if there are more samples/dialysis caps than can be accommodated on a single piece of membrane. Record the condition of each cap assembly in a notebook.
12. Attach the membrane/s with the dialysis chambers onto a Slide-A-Lyzer floatation assembly. Place the sample into the High-salt buffer.
13. Assemble the pump tubing to the Low-salt beaker, High-salt beaker, and Waste beaker, respectively (Figure 2B). Turn on the pump using a slow rate of 1.2 mL/min and let the pump run for 18 hours.
14. Increase the rate to 2.4 mL/min and run the pump until the Low-salt buffer is empty, which will take ∼ 45 minutes.
15. Add 1M DTT to the Zero-salt buffer and transfer the float assembly containing the membranes with the dialysis chambers into the Zero-salt buffer. Stir the buffer using a magnetic stir bar for at least 4 hours to complete the dialysis.
16. Take the membrane out of the Zero-salt buffer. Use a pipette to completely remove any buffer from the top of the dialysis chambers. Retrieve the nucleosome array assembly from inside the dialysis chambers by carefully puncturing the membrane with a pipette tip and pipetting the nucleosome solution into a new 0.5 mL tube. Store the nucleosome array assembly at 4 °C. The assembly is stable for a few months under this storage condition. Check the array occupancy using gel electrophoresis as described in section “Nucleosome assembly assayed by agarose gel” or perform a nucleosome array stretching assay as described in section “Stretching a nucleosome fiber with optical tweezers”.

**Figure 2:**
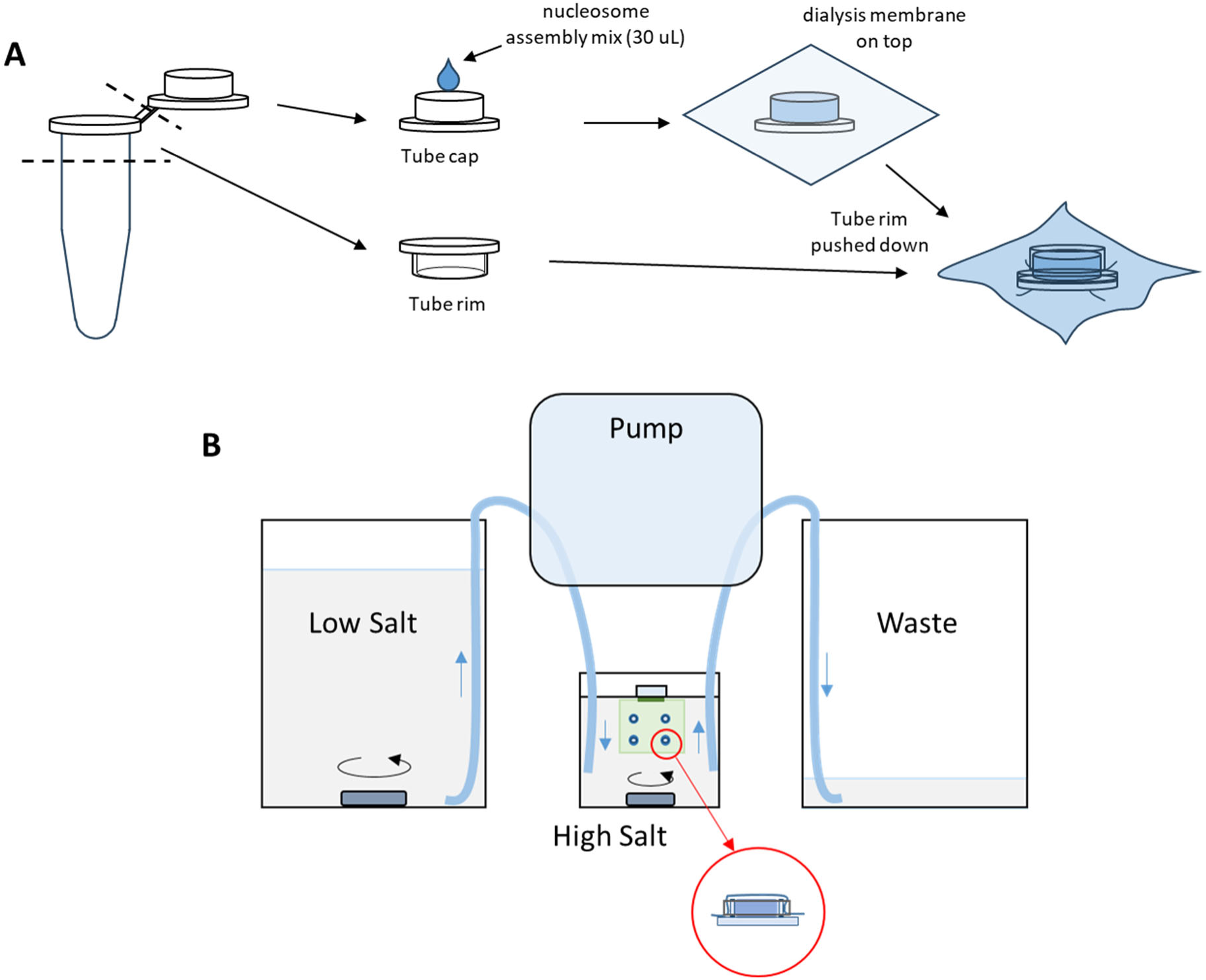
(A) Dialysis cap assembly. (B) Configuration of the continuous salt dialysis to assemble nucleosomes on the 64-mer 601 DNA template.

### Nucleosome assembly assayed by agarose gel

#### Timing: 1 hour

To quickly assess nucleosome occupancy at different molar ratios of histone octamers to 601 DNA repeats, perform native gel electrophoresis of the assembled chromatin as follows:

1. For agarose gel electrophoresis, dilute 2 µL of each nucleosome assembly in 8 µL of a Tris-EDTA buffer (10 mM Tris pH 8.0, 1 mM EDTA, 5% (v/v) glycerol) and load on a 10 cm, 0.7% agarose gel). Run the gel at 15 V/cm in 0.2X TBE (Tris-borate-EDTA) buffer for 30 minutes at room temperature and post-stain with ethidium bromide. As the assembly approaches saturation, the mobility of the high molecular weight band decreases and eventually plateaus, with concurrent formation of mono-nucleosomes assembled on the 147 bp competitor DNA (Figure 3).

**Figure 3:**
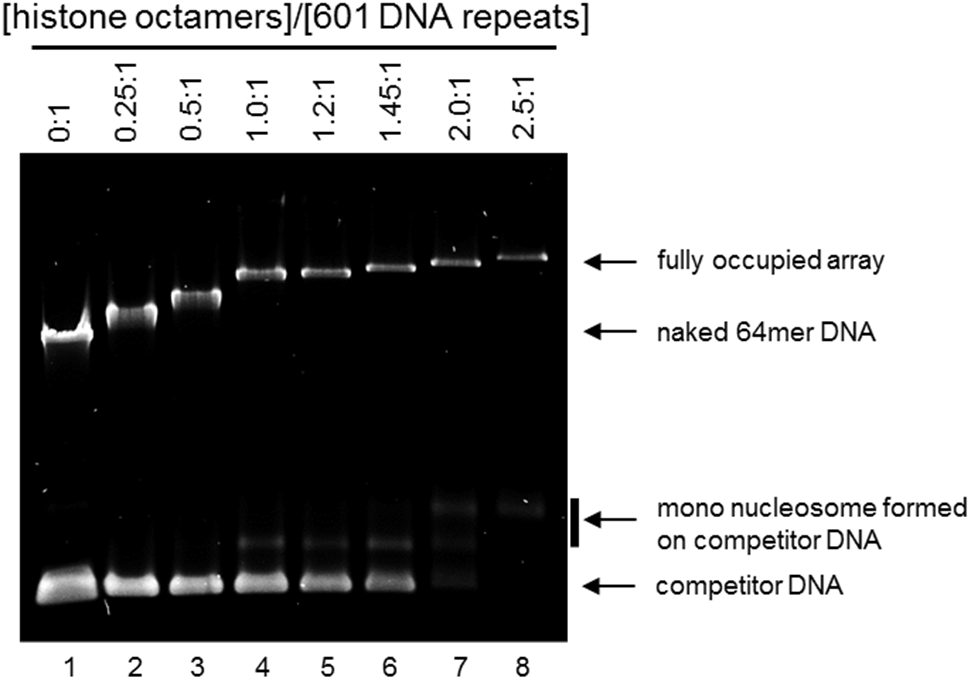
Native agarose electrophoresis of nucleosome arrays with increasing [histone octamers]:[601 DNA repeat] ratios. Figure reprinted and adapted with permission from Le et al., 2019^1^.

*Note:* Because nucleosomes limit ethidium bromide staining of the bound DNA, the saturation of a nucleosome assembly is more readily assessed by the disappearance of competitor DNA.

### Cleaning glass slides & coverslips

#### Timing: 1.5 hours

To remove visible dust and organic residues from the glass slides and glass coverslips, proceed with the following steps:

1. Spray Windex onto a glass slide and a glass coverslip. Use a folded Kimwipe to scrub the slide/coverslip several times and rinse them under running ultrapure H_2_0. Make sure all residue is washed away.
2. Place the cleaned slides and coverslips into separate clean Coplin jars. Fill the jars with ultrapure H_2_0 and sonicate them for 20 minutes.
3. Carefully discard the H_2_0 in the slide/coverslip holders and fill the jars with 95% EtOH. Sonicate for 20 minutes.
4. Carefully discard the EtOH. Rinse the slides/coverslips by filling the jars with fresh ultrapure H_2_0 and then discarding the H_2_0. Repeat this rinse three times. Then refill the holders with fresh ultrapure H_2_0 again and sonicate for 20 minutes.
5. Discard the H_2_0 a final time and fill the glass holders with fresh ultrapure H_2_0. Tightly cap the holders. Cleaned slides and coverslips can be stored in H_2_0 at room temperature until use.

### Nitrocellulose flow chamber preparation

#### Timing: 2 days

This method aims to produce a hydrophobic nitrocellulose-grease sample chamber. Nitrocellulose increases the hydrophobicity of a glass surface, which can be further enhanced by high vacuum grease. Slow diffusion of the vacuum grease deposits dimethyl-and polymethyl-siloxane groups over the nitrocellulose-coated surface. The increased surface hydrophobicity, when combined with a purified blocking protein beta-casein, results in an improved surface passivation capacity against nonspecific surface adsorption of nucleosome fibers and topoisomerases.

1. Take a cleaned glass slide and coverslip as described in the section “Cleaning glass slides & coverslips” and dry completely under an air flow.
2. Put the slide in a clean empty tip box. Drop ∼ 70-80 µL of 1-2% nitrocellulose solution onto the slide. See “Materials and Equipment” for nitrocellulose solution preparation (Figure 4).
3. Gently drop a clean coverslip over the slide and allow the nitrocellulose solution to spread over the entire surface between the slide and the coverslip.

**Figure 4.**
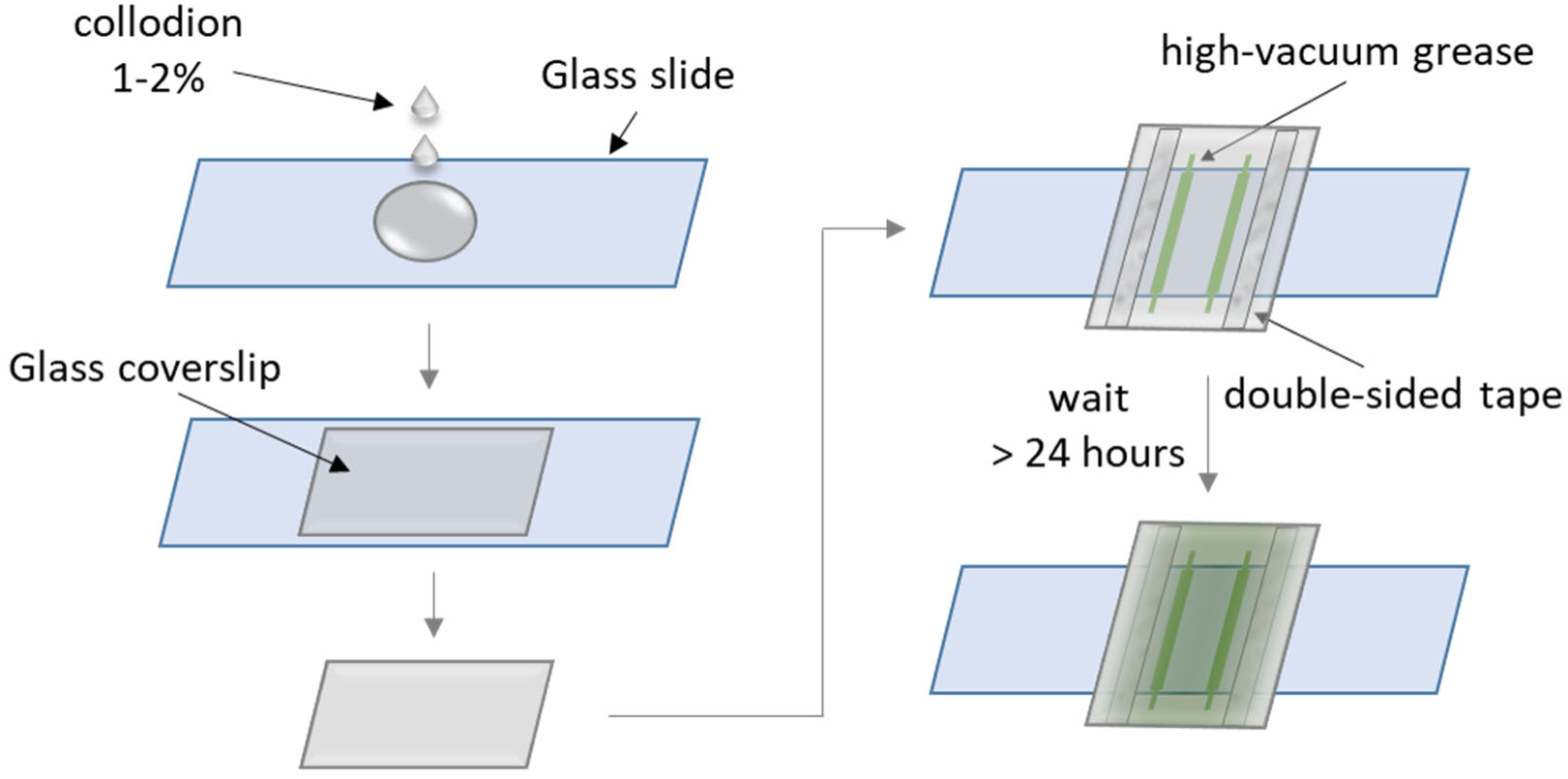
Steps to create the hydrophobic nitrocellulose-coated grease sample chamber MT stuck bead fiducial marker preparation

*Note:* Avoid introducing large bubbles. Small bubbles will typically move to the edge of the coverslip due to capillary force.

4. Use a clean pair of tweezers to carefully separate the slide and coverslip.

*Note:* Do not lift the coverslip off the slide but slide the coverslip off the slide to obtain a uniform nitrocellulose coating on the coverslip’s surface.

5. Gently dry the slide and the coverslip under an airflow.
6. Gently drop the coated coverslip into a 50-mL tube filled with 95% EtOH and incubate at room temperature for 5 minutes.

*CRITICAL:* 95% EtOH will further harden the nitrocellulose coat. Do not use 100% EtOH, as it may crack the nitrocellulose layer.

7. While waiting for the coverslip, place the slide with the nitrocellulose coat under running distilled H_2_0 to remove the nitrocellulose film on the slide. Dry the slide under an air flow and let it sit in a clean box.

*Note:* This step recycles the slide used in nitrocellulose coating. A freshly cleaned slide can also be used each time.

8. Decant the EtOH and dry the coated coverslip in the empty tube in an oven at 80 °C for 10 minutes. Then let the coated coverslip cool to room temperature.
9. Rinse the coverslip by very gently filling the tube that holds the coated coverslip with ultrapure H_2_0. Slowly decant the H_2_0. Repeat this step one time.

*CRITICAL:* Perform carefully as a strong water flow can easily strip off the nitrocellulose film.

10. Dry the coverslip using an airflow. Assemble a flow chamber using one cleaned slide and the coated coverslip. Use high-vacuum grease to create two parallel walls for the flow chamber as shown in Figure 4. Then position the slide on top of the grease and firmly press down on the taped areas to secure the sample chamber.

*Note:* Use a blunt tip syringe pre-filled with vacuum grease to conveniently dispense the grease onto the coverslip.

11. Keep the sample chamber in a clean tip box for more than 24 hours at room temperature before using so that the flow chamber becomes hydrophobic due to grease spreading over the chamber surface.

*Note:* For consistency in sample chamber performance, it is preferred to keep the sample chamber in a wet tip box (a tip box containing ultrapure H_2_0 that does not touch or cover the slide) for 2 days before use. Incubation longer than two days leads to a significant degree of hydrophobicity that can prevent solutions from being pulled into the chamber via capillary force so that the chamber cannot be used for experiments.

### MT stuck bead fiducial marker preparation

#### Timing: 1 hour

After preparing the surface-anchored nucleosome sample as will be described in Section F, the sample chamber is mounted onto an optical tweezers or a magnetic tweezers instrument. To obtain accurate extension measurements on magnetic tweezers and correct for instrument drift, it is essential to generate fiducial markers. In previous experiments, a small population of magnetic beads introduced during the tether anchoring step would nonspecifically get stuck on the surface and those stuck beads were employed as fiducial markers. However, the improved surface passivation of our described protocol effectively suppresses nonspecific bead sticking. Therefore, we needed to manually generate beads that are stuck on the surface that serve as fiduciary markers. To generate stable fiducial markers under our surface conditions, we developed a method using biotin-and-digoxygenin-labeled DNA to strongly anchor streptavidin-coated magnetic beads to an anti-digoxygenin-coated surface that could resist magnetic pulling under a few tens of piconewton forces.

*Note:*

1. Make the double-labeled sticky DNA adapters (labeled with both biotin and digoxygenin) as described in “Materials and Equipment”
2. Prepare the double-labeled 500-bp DNA-coated magnetic beads

a. In a 500-µL tube, add 0.35 µL of washed streptavidin-coated T1 magnetic beads into 99 µL PBS 1X. Mix and sonicate for 30 seconds. For optimal performance, wash the beads as described in the section Wash the streptavidin-coated magnetic beads or streptavidin-coated cylinders.
*CRITICAL:* It is critical to use the Dynabeads MyOne Streptavidin T1 magnetic bead when working with nucleosome fibers. We found that other streptavidin-coated magnetic beads, such as Dynabeads MyOne Streptavidin C1 from Thermo Fisher or S1420S from NEB, could induce histone loss, potentially due to the iron leakage from the bead core in a buffer containing EDTA.

b. Cool the bead solution down by leaving the tube in ice for 40 seconds. Then add 2.5 µL of the 93-nM 500-bp biotin-dig-labeled adapter. Mix well and incubate at 4 °C for 30 minutes.
c. Pull down the magnetic beads by putting the tube in a magnet rack. Discard the supernatant and then add 50 µL PBS 1X and re-suspend the beads. Store this bead solution at 4 °C.
*CRITICAL:* It is critical to discard the supernatant with unbound sticky DNA adapters as it could be readily adsorbed to the surface and compete with DNA tethering during the DNA anchoring steps.
3. Generate the stuck bead fiducial marker

a. Incubate a hydrophobic nitrocellulose-coated grease sample chamber with ∼ 50 µL of 1-20 ng/µL anti-digoxygenin for 30 minutes
b. Dilute 2.5 µL of the sticky DNA-bound bead solution, as prepared in the previous steps, into 65 µL of PBS 1X and mix well. Sonicate for 10 seconds.
c. Flow the mixture into the sample chamber and incubate for 10 minutes at room temperature.
d. Flush out the unbound beads by introducing 95 µL beta-casein 25 mg/mL (see “Beta-casein preparation”) to start the surface passivation step.
e. On a Magnetic Tweezers (MT), mount the sample chamber and check for stuck beads. Following the described protocol, there are typically ∼ 10 stuck beads per a 562 µm x 350 µm FOV. Quickly rotate the magnet to identify stably stuck beads (which stay fixed under magnetic rotation) and use these beads as references.

*Note:* If there are too few or too many stuck beads, increase or decrease the amount of DNA-bound beads in the dilution in Step 3, respectively.

### Beta-casein preparation

#### Timing: 1 day

Beta-casein is a phosphoprotein and is one of four types of milk proteins found in mammals (αS1, αS2, β, and κ). While it has previously been used to suppress bead sticking in experiments with motor proteins^19^, beta-casein has not been used for single-molecule studies with chromatin. We found that beta-casein is highly effective in reducing non-specific surface sticking of large chromatin tethers in physiological buffer conditions containing magnesium at millimolar levels^1^. 40 mL beta-casein solution from bovine milk at 25 mg/mL is prepared using the following protocol:

1. Add 400 µL 1M Tris-Cl pH 8.3 directly into the beta-casein bottle. Shake the bottle a few times.
*Note:* Use a basic solution to compensate for the mild decrease in the solution pH upon casein dissolution.
2. Add 400 µL 5 M NaCl and then add 30 mL ultrapure H_2_O into the same bottle. Tightly cap the bottle and use parafilm to securely seal the cap.
3. Firmly affix the bottle onto a vertical wheel using strong tape. Spin the wheel at 20% power overnight at 4 °C.
4. The next day, take the beta-casein bottle off the wheel and incubate at 4 °C for 2 hours until all the bubbles disappear.
5. Aliquot the beta-casein solution into 1.5 mL tubes and store at -80 °C for long-term storage.
6. For short-term storage, thaw an aliquot at 4 °C, pipette up and down 7 times to mix, centrifuge for 5 seconds, and aliquot into smaller volume tubes. Store aliquots at -20 °C.

### Wash the streptavidin-coated magnetic beads or streptavidin-coated cylinders

#### Timing: 30 minutes

To obtain optimal tether density, it is recommended to wash the streptavidin-coated magnetic beads or cylinders, preferably before each experiment. We noted that several week-old bead/cylinder samples have reduced tether anchoring efficiency, potentially due to bound streptavidin getting released into the bead solution and hence, the free streptavidin may compete for tether binding.

1. For streptavidin-coated cylinder solution, spin down at 5000 rpm for 10 minutes. For streptavidin magnetic-coated bead solution, pull down the beads with a magnetic rack (Thermo Fisher, FERMR01). Carefully pipette and discard the supernatant.
2. Add an equal volume of PBS 1X and resuspend the beads/cylinders by pipetting.
3. Repeat steps 1 and 2.
4. Repeat step 1 and replace with an equal volume of bead storage buffer containing PBS 1X and 1 mg/mL BSA. Store the bead/cylinder sample at 4 °C.

*Critical:* Do not freeze the bead/cylinder solution.

## D. KEY RESOURCE TABLE

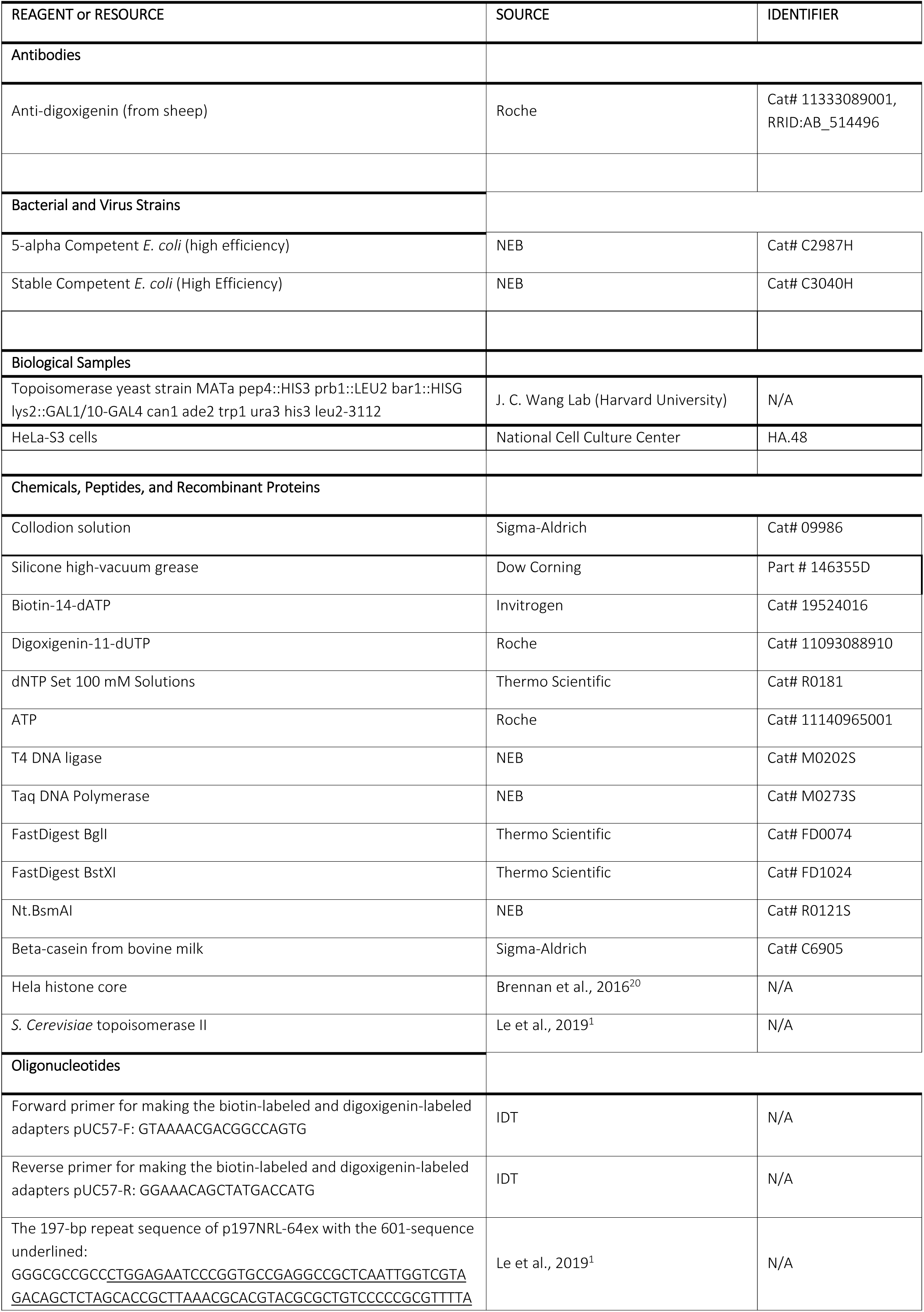
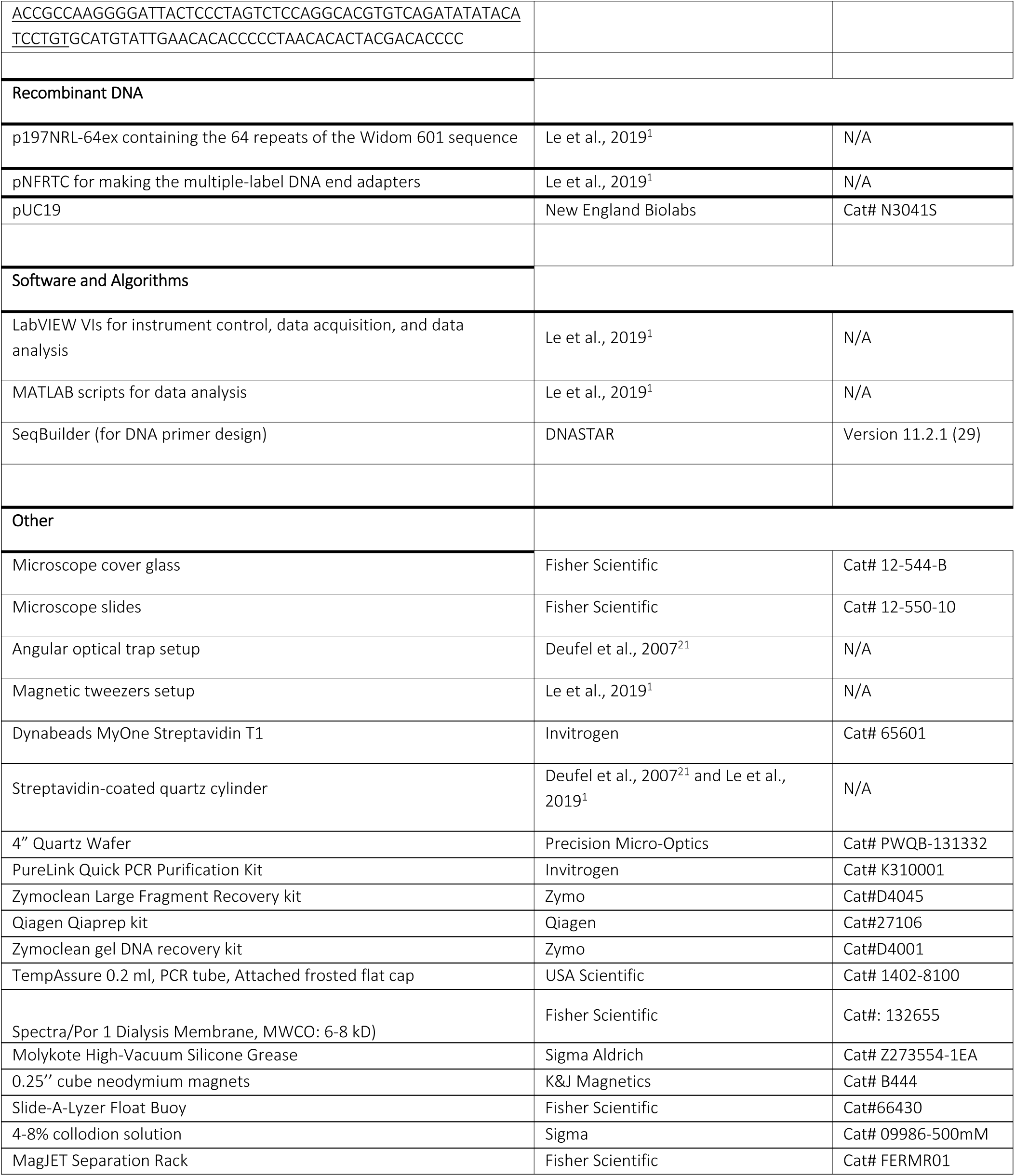

## E. MATERIALS AND EQUIPMENT SETUP

### Angular optical tweezers

By rotating the trapping beam’s linear polarization, our angular optical tweezers (AOT) allow simultaneous control and measurement of rotation, torque, displacement, and force of a trapped nanofabricated quartz cylinder about its cylindrical axis^21–28^.

### Magnetic tweezers

Our custom-built MT setup^1,12,13^ was based on previous designs^29,30^. The magnetic field was generated with a pair of 0.25’’ cube neodymium magnets which were arranged with their dipoles oriented in opposing directions and parallel to the optical axis of the microscope, with a separation gap of 0.5 mm. Magnetic bead images were collected using a Nikon 40x objective lens (Plan Apo40x 0.95 NA) on a 2.3 MP camera (Basler acA1920-155um) at a frame rate of 10 fps and an exposure time of 0.5 ms. The bead positions were tracked in three dimensions using an algorithm implemented in LabVIEW based on the source code available on Omar Saleh’s website^31^.

### Preparation of solutions

#### (1) 1-2% Nitrocellulose Solution

| Reagent | Final Concentration | Amount |
| --- | --- | --- |
| 4-8% collodion solution (Cat# 09986-500mM, Sigma) | 1-2% | 10 mL |
| Amyl Acetate |  | 30 mL |
| Total |  | 40 mL |

In a fume hood, add reagents to a 50 mL falcon tube and invert the tube a few times to mix. Tightly close the tube cap to prevent evaporation of the solvent.

#### (2) NaN_3_ 20 mg/mL Solution

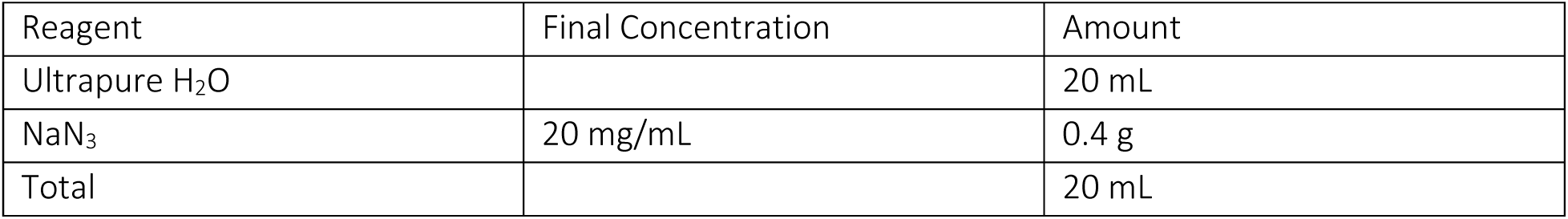

Add reagents to a 50 mL falcon tube and invert the tube a few times to mix. NaN_3_ is neurotoxic. Wear gloves and a face mask when handling the powder.

#### (3) Nucleosome Dialysis Buffers

Low-Salt Buffer

| Reagent | Final Concentration | Amount |
| --- | --- | --- |
| Ultrapure H<sub>2</sub>O |  | 1297.8 mL |
| Tris-Cl 1M pH 8.0 | 10 mM | 14 mL |
| EDTA 0.5 M | 1 mM | 2.8 mL |
| NaCl 5 M | 250 mM | 70 mL |
| DTT 1M | 1 mM | 1.4 mL |
| NaN<sub>3</sub> 20 mg/mL | 0.2 mg/mL | 14 mL |
| Total |  | 1400 mL |

High-Salt Buffer

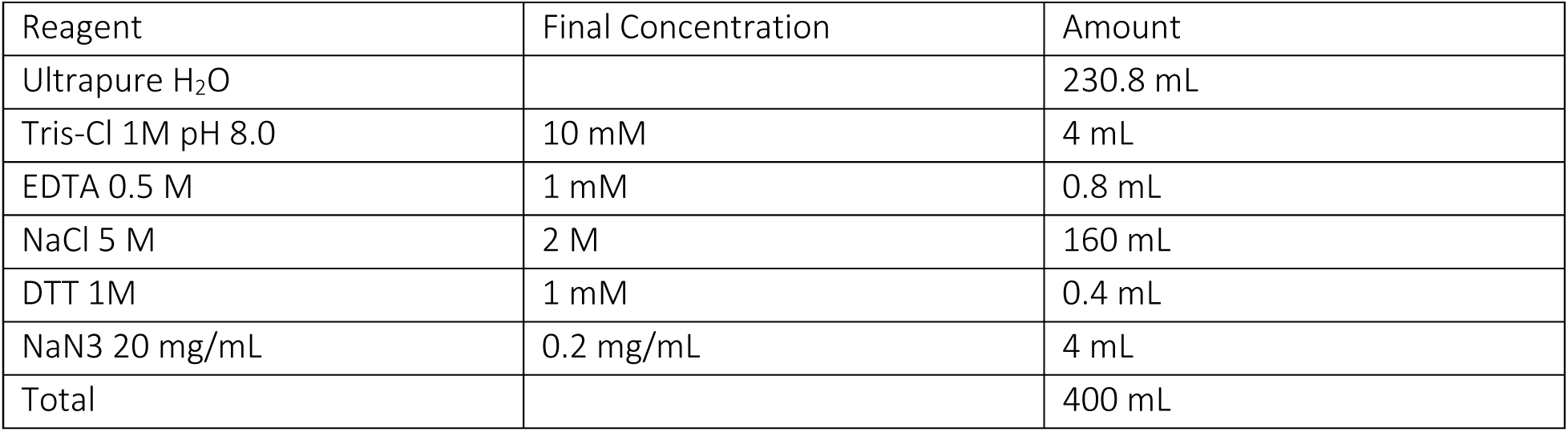

| Reagent | Final Concentration | Amount |
| --- | --- | --- |
| Ultrapure H<sub>2</sub>O |  | 230.8 mL |
| Tris-Cl 1M pH 8.0 | 10 mM | 4 mL |
| EDTA 0.5 M | 1 mM | 0.8 mL |
| NaCl 5 M | 2 M | 160 mL |
| DTT 1M | 1 mM | 0.4 mL |
| NaN<sub>3</sub> 20 mg/mL | 0.2 mg/mL | 4 mL |
| Total |  | 400 mL |

Zero-Salt Buffer

| Reagent | Final Concentration | Amount |
| --- | --- | --- |
| Ultrapure H<sub>2</sub>O |  | 394.8 mL |
| Tris-Cl 1M pH 8.0 | 10 mM | 4 mL |
| EDTA 0.5 M | 1 mM | 0.8 mL |
| DTT 1M | 1 mM | 0.4 mL |
| Total |  | 400 mL |

The Zero-salt and High-salt buffers are kept in 500-mL beakers and the Low Salt buffer is kept in a 2000-mL beaker. Use an autoclaved 100-mL graduated cylinder for transferring solutions (i.e., ultrapure H_2_O, NaCl 5M). Thaw and add DTT to the buffer immediately before use to prevent degradation.

#### (4) Nucleosome Assembly 2X Buffer

| Reagent | Final Concentration | Amount |
| --- | --- | --- |
| Ultrapure H<sub>2</sub>O |  | 252 µL |
| Tris-Cl 1M pH 7.5 | 40 mM | 80 µL |
| EDTA 0.5 M | 2 mM | 8 µL |
| NaCl 5 M | 4 M | 1.6 mL |
| DTT 1M | 20 mM | 40 µL |
| acetylated BSA 20 mg/mL | 0.2 mg/mL | 20 µL |
| Total |  | 2000 µL |

Store the nucleosome assembly buffer at -20 °C.

#### (5) Chromatin Dilution 10X Buffer

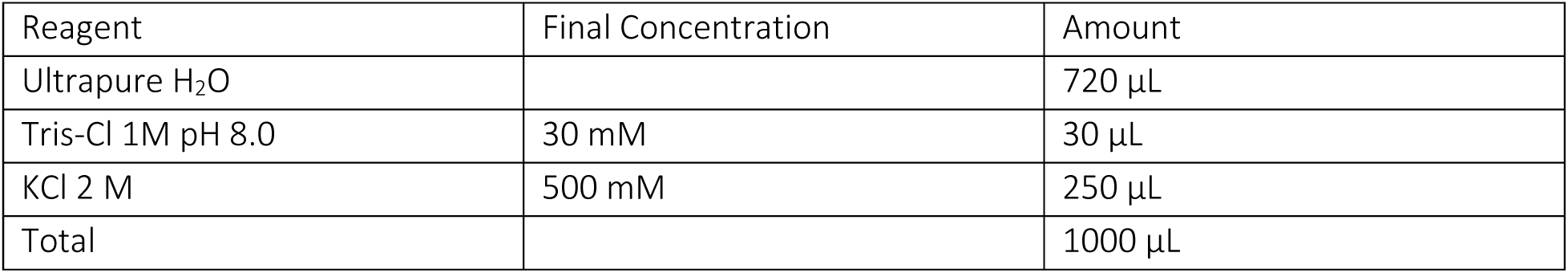

| Reagent | Final Concentration | Amount |
| --- | --- | --- |
| Ultrapure H<sub>2</sub>O |  | 720 µL |
| Tris-Cl 1M pH 8.0 | 30 mM | 30 µL |
| KCl 2 M | 500 mM | 250 µL |
| Total |  | 1000 µL |

#### (6) Topoisomerase Dilution Buffer

| Reagent | Final Concentration | Amount |
| --- | --- | --- |
| Ultrapure H<sub>2</sub>O |  | 715.5 µL |
| TCEP 1 M | 0.5 mM | 0.5 µL |
| beta-casein 25 mg/mL | 0.1 mg/mL | 4 µL |
| Total |  | 1000 µL |

Store the topoisomerase dilution buffer at -20 °C. Add beta-casein immediately before use.

#### (7) Topoisomerase Reaction Buffer

|  | Final Concentration | Amount |
| --- | --- | --- |
| Ultrapure H<sub>2</sub>O |  | 1760.6 µL |
| Tris-Cl 1M pH 8.0 | 10 mM | 20 µL |
| NaCl 5 M | 50 mM | 20 µL |
| KCl 2 M | 50 mM | 50 µL |
| MgCl<sub>2</sub> 1 M | 3 mM | 6 µL |
| EDTA 500 mM | 0.1 mM | 0.4 µL |
| ATP 100 mM | 1 mM | 20 µL |
| DTT 1 M | 1 mM | 2 µL |
| TCEP 1 M | 0.5 mM | 1 µL |
| beta-casein 25 mg/mL | 1.5 mg/mL | 120 µL |
| Total |  | 2000 µL |

#### (8) Topoisomerase Flushing Buffer

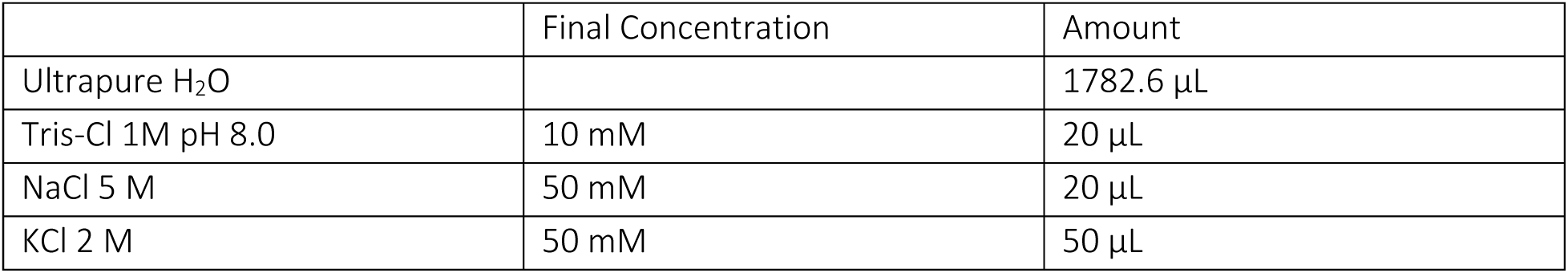
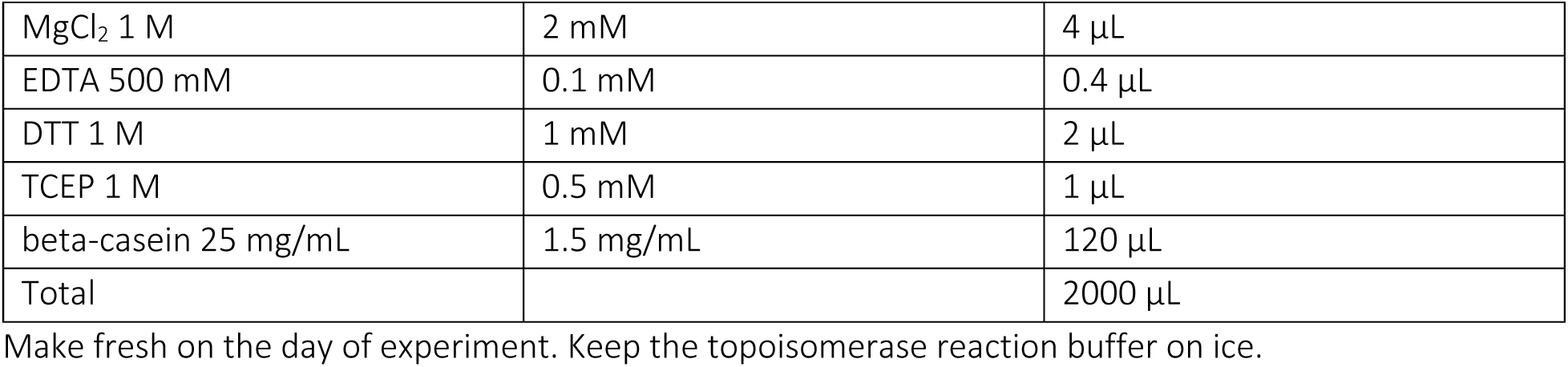

|  | Final Concentration | Amount |
| --- | --- | --- |
| Ultrapure H<sub>2</sub>O |  | 1782.6 µL |
| Tris-Cl 1M pH 8.0 | 10 mM | 20 µL |
| NaCl 5 M | 50 mM | 20 µL |
| KCl 2 M | 50 mM | 50 µL |
| MgCl<sub>2</sub> 1 M | 2 mM | 4 µL |
| EDTA 500 mM | 0.1 mM | 0.4 µL |
| DTT 1 M | 1 mM | 2 µL |
| TCEP 1 M | 0.5 mM | 1 µL |
| beta-casein 25 mg/mL | 1.5 mg/mL | 120 µL |
| Total |  | 2000 µL |

Make fresh on the day of experiment. Keep the topoisomerase reaction buffer on ice.

### Preparation of DNA templates

#### (1) Biotin-labeled/Dig-labeled 500-bp DNA adapter

- Mix reagents in PCR tubes precooled on ice. Use separate tubes for each unique adapter. Add DNA polymerase last. Keep the tubes on ice after mixing. Both adapters are needed to achieve the desired template. pNFRTC (or pMDW111) is a plasmid that contains a low nucleosome affinity sequence used for making the sticky adapters for the torsionally-constrained template.

25%-biotin-labeled adapter

| Reagent | Final Concentration | Amount |
| --- | --- | --- |
| Ultrapure H<sub>2</sub>O |  | 136.3 µL |
| Forward primer 100 µM | 0.5 µM | 0.8 µL |
| Reverse primer 100 µM | 0.5 µM | 0.8 µL |
| pNFRTC (pMDW111) 1 ng/µL | 2 ng/50 µL | 8.0 µL |
| dNTPs 25 mM | 0.2 mM | 1.6 µL |
| biotin dATP 0.4 mM | 0.067 mM | 33.5 µL |
| Taq buffer 10X | 1X | 20.0 µL |
| Taq DNAP | 0.5 unit/50 µL | 1.0 µL |
| Total |  | 200 µL |

25%-digoxygenin-labeled adapter

25%-digoxigenin-labeled adapter
| Reagent | Final Concentration | Amount |
| --- | --- | --- |
| Ultrapure H<sub>2</sub>O |  | 156.4 µL |
| Forward primer 100 µM | 0.5 µM | 0.8 µL |
| Reverse primer 100 µM | 0.5 µM | 0.8 µL |
| pNFRTC (pMDW111) 1 ng/µL | 2 ng/50 µL | 8.0 µL |
| dNTPs 25 mM | 0.2 mM | 1.6 µL |
| dig dUTP 1 mM | 0.067 mM | 13.4 µL |
| Taq buffer 10X | 1X | 20.0 µL |
| Taq DNAP | 0.5 unit/50 µL | 1.0 µL |
| Total |  | 200 µL |

- Raise the temperature of the heat block to 95 °C and place the PCR tubes into the heat block before running the following heat cycle. Prepare the reagents for 4 PCR reactions for each adapter and divide equally into 4 0.2-mL PCR tubes.

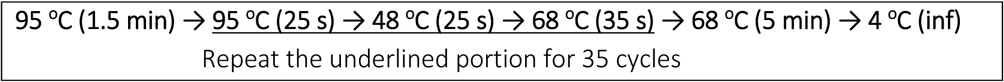

- After PCR, check the DNA on a 10 cm, 1% agarose gel. Purify DNA using Pure Link PCR spin columns. DNA is eluted using an elution buffer (10 mM Tris-Cl pH 8.0 and 0.1 mM EDTA) prewarmed at 50 °C. Store the DNA at 4 °C.

#### (2) 147-bp DNA competitor

PCR amplify the 147-bp DNA competitor:

- Mix the following ingredients in a PCR tube precooled on ice. Add DNA polymerase last. Keep the tube on ice after mixing.

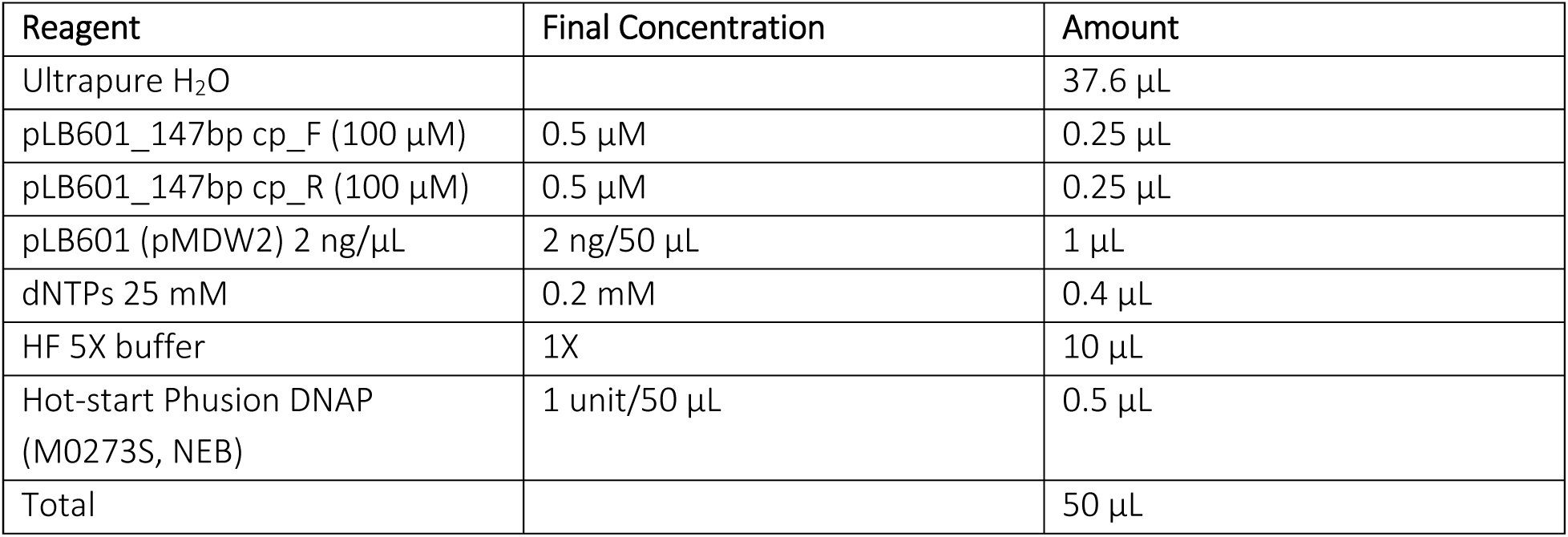

| Reagent | Final Concentration | Amount |
| --- | --- | --- |
| Ultrapure H<sub>2</sub>O |  | 37.6 µL |
| pLB601_147bp cp_F (100 µM) | 0.5 µM | 0.25 µL |
| pLB601_147bp cp_R (100 µM) | 0.5 µM | 0.25 µL |
| pLB601 (pMDW2) 2 ng/µL | 2 ng/50 µL | 1 µL |
| dNTPs 25 mM | 0.2 mM | 0.4 µL |
| HF 5X buffer | 1X | 10 µL |
| Hot-start Phusion DNAP (M0273S, NEB) | 1 unit/50 µL | 0.5 µL |
| Total |  | 50 µL |

- Raise the temperature of the heat block to 98 °C and place the PCR tubes into the heat block before running the following heat cycle. Prepare the reagents for 20 PCR reactions and divide equally into twenty 0.2-mL PCR tubes.

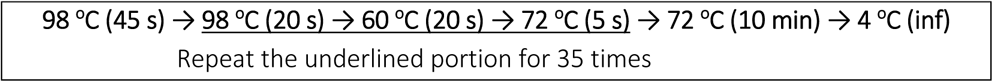

- After PCR, check the DNA on a 10 cm, 1% agarose gel. Purify DNA using Pure Link PCR spin columns. Store the DNA at 4 °C.

#### (3) Biotin-digoxygenin-labeled ‘sticky’ DNA adapter

- Mix the following ingredients in PCR tubes precooled on ice. Add DNA polymerase last. Keep the tubes on ice after mixing. pNFRTC (or pMDW111) is a plasmid that contains a low nucleosome affinity sequence used for making the sticky adapters for the torsionally-constrained template.

| Reagent | Final Concentration | Amount |
| --- | --- | --- |
| Ultrapure H<sub>2</sub>O | | 79.7 μL |
| Forward primer (100 μM) | 0.4 μM | 0.48 μL |
| Reverse primer (100 μM) | 0.4 μM | 0.48 μL |
| pNFRTC (pMDW111) 1 ng/μL | 2 ng/50 μL | 4.8 μL |
| dNTPs 25 mM | 0.2 mM | 0.96 μL |
| biotin dATP 0.4 mM | 0.05 mM | 15 μL |
| dig dUTP 1 mM | 0.05 mM | 6 μL |
| Taq buffer 10X | 1X | 12 μL |
| Taq DNAP<br>(M0273S, NEB) | ~ 1 unit/50 μL | 0.6 μL |
| Total | | 120 μL |

- PCR amplify using the following heating/cooling cycle. Raise the temperature of the heat block to 95 °C and place the PCR tubes into the heat block before running the following heat cycle.

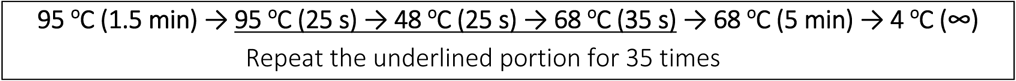

- After PCR, purify DNA using Pure Link PCR spin column. DNA is eluted using an elution buffer (10 mM Tris-Cl pH 8.0 and 1 mM EDTA) prewarmed at 50 °C. Check the DNA concentration using a spectrophotometer.

## F. STEP-BY-STEP METHOD DETAILS

### Forming single fiber tethers with nucleosome arrays for optical tweezers or magnetic tweezers

#### Timing: 2-4 hours

This protocol was designed to form chromatin tethers with ∼ 50 nucleosomes assembled on the 64-mer DNA template. Use sample chamber construction and nucleosome assembly protocols detailed in “BEFORE YOU BEGIN”.

1. Incubate a hydrophobic nitrocellulose-coated grease sample chamber (see “BEFORE YOU BEGIN”) with 20 ng/µL anti-digoxygenin in PBS 1X and incubate for 30 minutes.
2. If doing an MT experiment, you will need to generate fiducial markers. To do this, flow in the beads coated with the sticky adapters (see “BEFORE YOU BEGIN”) and wait for 10 minutes. If doing an AOT experiment, skip this step.
3. Flow in 90 µL of 25 mg/mL beta-casein (see “BEFORE YOU BEGIN”) and incubate for 1-3 hours at room temperature.
4. Flow in ∼ 50-100 µL of 20 pM chromatin fibers diluted in CD 1X buffer (diluted from the chromatin dilution (CD) 10X buffer, see “Materials and Equipment”) with 1.5 mg/mL beta-casein added and incubate for 15-20 minutes at room temperature.
5. Flow in ∼ 90 µL of streptavidin-coated polystyrene beads or streptavidin-coated magnetic bead or streptavidin-coated quartz cylinders at ∼ 1 pM diluted in CD 1X buffer. Incubate for 15-20 minutes to allow anchoring of the bead or cylinder to the chromatin tether.

*Note:* To remove bead/cylinder clustering, sonicate the bead/cylinder solution for 30 seconds and cool the bead solution on ice for 10 seconds before flowing into the sample chamber

Flush the sample chamber with 100 µL of CD 1X buffer to remove free beads/cylinders.

6. Flow in 75-100 µL of the working buffer. An example is the topo reaction buffer (see “Materials and Equipment”).
7. Proceed immediately to data acquisition on optical tweezers or magnetic tweezers

*Note:* Following this preparation, the sample is stable for data acquisition for 2-3 hours without noticeable increase in bead sticking over time. If the tether density is low, re-wash the cylinder/bead solution using the protocol described in “BEFORE YOU BEGIN”.

### Twisting and stretching nucleosome fibers with (angular) optical tweezers

#### Timing: 1-3 hours

1. Mount the sample chamber with the anchored chromatin tethers on the optical tweezers set-up. Position the stage so that the imaging plane is slightly above the bead/cylinder.
2. For twisting a chromatin fiber on the AOT, stretch the tether vertically to 0.5 pN, and introduce turns with a rate of 4 turns/s. Typically, to obtain continuous winding curves in the forward and the reverse directions for a 50-nucleosome array, wind the tether over a range between -35 and +70 turns, from turn 0 → -35 → 0 → +70 → 0 → -35 → 0. The winding curves are obtained between turn -35 and turn +70.
3. For stretching a chromatin fiber, use a velocity clamp with a rate of 200-400 nm/s to disrupt the histone octamers
4. Move to a new tether within the sample chamber and repeat the process.

Continuous winding experiments of single fibers on the MT Timing: 30 minutes

1. Mount the sample chamber with the anchored chromatin tethers on the magnetic tweezers set-up.
2. Select tethered beads and set the magnet height to obtain 0.5 pN on average (*h*_magnet_ ∼ 4.7 mm). Set the acquisition time to be 50 ms and the frame rate to be 10 Hz.
3. Construct a look-up table for the height measurement, determine the tether attachment point, and measure the Z-tension from the lateral fluctuations of the magnetic bead.
4. Plot an initial winding curve by rotating the magnets between turn -30 and turn +37 at a rate of 10 turns/s. At each integer turn number, hold the magnet still for 500 ms. Return to turn 0.
5. Dilute *S. cerevisiae* topoisomerase II (scTopo II) to specified concentration using a 3-step protocol as described below:

a. Dilute topo II from 2 uM to 50 nM by adding 4 µL of the 2-uM stock topo II into 156 µL of a topo dilution buffer (see “Materials and Equipment”). This dilution can be kept on ice and used for further dilutions for 2-3 hours.
b. Immediately before the experiment, dilute topo II from 50 nM to 0.25 – 1 nM in topo dilution buffer.
c. Further dilute topo II from 0.25 – 1 nM to 1.5 – 10 pM in the topo reaction buffer. This is the topo sample to be introduced into the sample chamber.

*Note:* To maintain the consistency in pipetting of a solution with glycerol, avoid pipetting < 0.5-µL volume when diluting topo II. For consistency in diluting the topo II sample, carefully mix the solution by slowly but consistently pipetting up and down 70% of the total volume 7-8 times.

6. Flow ∼ 95 µL of the diluted topo solution into the sample chamber and incubate for 2 minutes to achieve equilibration in topo II binding to the chromatin tether.
7. Set the magnet rate to 3.6 turns/s and introduce +1000 turns under this winding rate.
8. After the magnet stops, flush the sample chamber with 95 µL of the topo flushing buffer (see Materials and Equipment) to remove unbound topo and deactivate topo activity.
9. Plot the final winding curve from turn +970 to +1100 under a winding rate of 10 turns/s while holding for 500 ms at each integer turn number. At the end of the protocol, torsionally constrained tethers are wound to the surface. This information is used to determine the absolute z-extension of the tethers.

## G. EXPECTED OUTCOMES

### Stretching a nucleosome fiber with optical tweezers

The optical tweezers data is converted into extension and force using a LabView-based data conversion software. Figure 5 represents the stretching data of ∼ 50 nucleosome fiber before and after surface passivation optimization. Without proper surface passivation, most of the fiber tethers were short, and their force-extension data revealed irregular saw-toothed behavior with frequent big force drops, indicative of continuous pulling of a partially stuck fiber off the surface (pink curve in Figure 5). With our efficient surface blocking method, ∼ 90% of the stretching curves showed consistent data of unwrapping an intact nucleosome fiber: unwrapping starts first with the gradual outer-turn release of nuclesomes under low force (< 10 pN), followed by the uniform inner-turn release of nucleosomes at ∼ 25 pN force, until the fiber is fully unraveled with the stretching curve mimicking that of naked DNA at >= 40 pN forces (green curve in Figure 5). This behavior is fully consistent with previous reports on stretching a smaller nucleosome array^3,5^.

**Figure 5:**
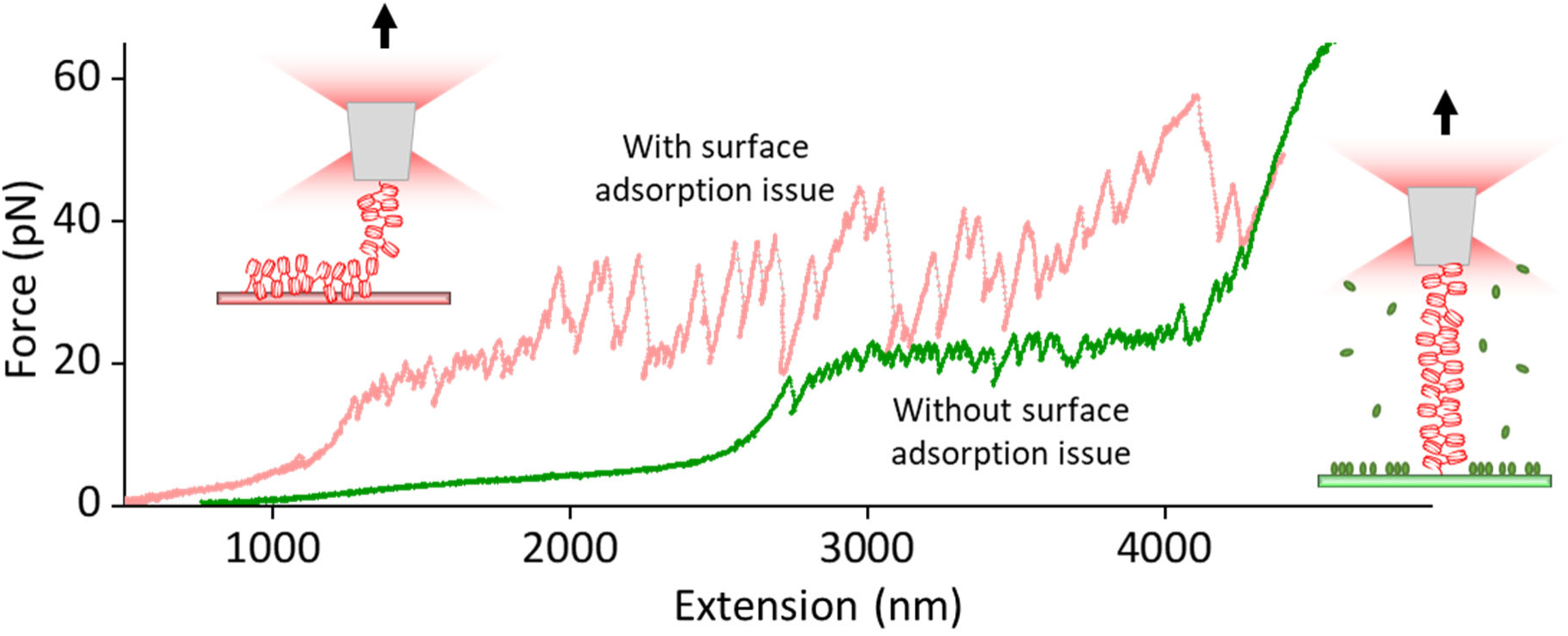
Representative force-extension data of stretching a nucleosome fiber under poorly passivated surface (pink curve) and properly passivated surface using the protocol described in this paper (green curve).

### Twisting and stretching a nucleosome fiber with angular optical tweezers

The angular optical tweezers data is converted into extension, force, and torque using a LabView-based data conversion software. Figure 6A represents typical twisting data with tether extension vs. number of turns added/removed for a chromatin fiber under a constant tension of 0.5 pN. The minimal difference seen between the forward and reverse winding curves indicates the twisting process is in equilibrium without histone dissociation. The winding curve is essentially asymmetric, wider on positive turns, which suggests that a single chromatin fiber is capable of absorbing more positive twists than negative twists without significant collapse. Figure 6B represents a subsequent stretching curve of the same chromatin fiber without any surface sticking. We used the stretching data to determine the quality and number of nucleosomes in the single fiber.

**Figure 6:**
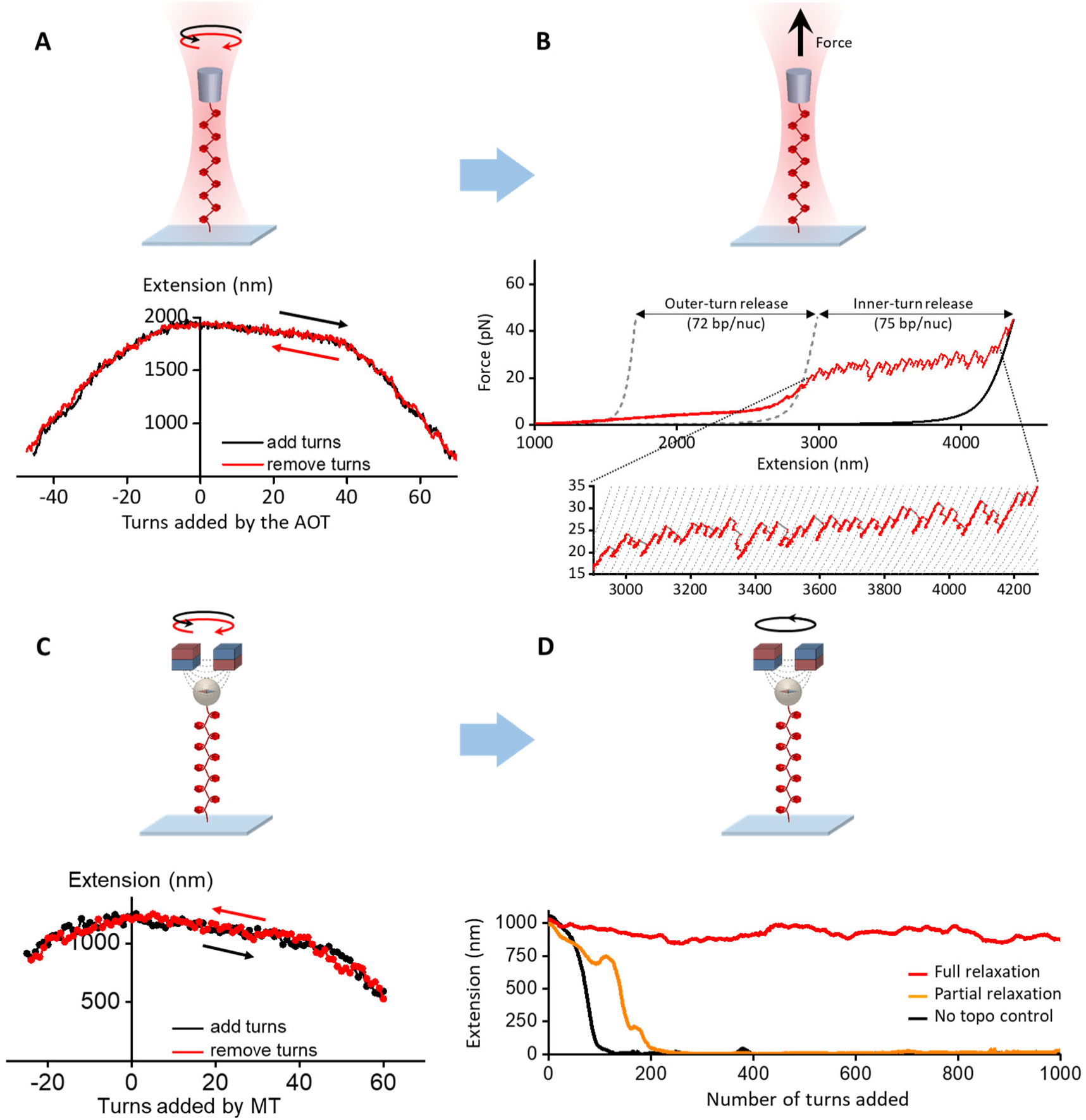
(A) Single chromatin fiber stability under twisting using an angular optical trap. Shown is an example trace of adding (black) and removing (red) turns, resulting in two hat curves. (B) An example trace of the force-extension curve of a single chromatin fiber containing ∼ 50 nucleosomes after the initial twisting, showing 72 bp of smooth outer-turn DNA release and 75 bp of sudden inner-turn release from each nucleosome. Outer-turn DNA release starts at 2 pN and ends at 15 pN, before the inner-turn DNA release starts, similar to what we have reported previously^3,5^. The two gray dashed curves^32^ correspond to naked DNA of lengths such that their force-extension curves cross the chromatin fiber curve at 2 pN and 15 pN and they are used to characterize the amount of outer-turn DNA released. The black solid curve corresponds to naked DNA whose number of base pairs is the same as that of the chromatin fiber’s DNA (12,667 bp). The dashed curves of the inset are naked DNA curves with 75 bp increments in length. (C) Single nucleosome fiber under twisting using magnetic tweezers. Shown is an example trace of adding (black) and removing (red) turns, resulting in two hat curves. (D) Example traces of single chromatin fiber extension during the continuous winding step in the presence of topoisomerase II using magnetic tweezers. The magnetic bead added 1000 winding turns at 3.6 turns/s. By removing the added turns, topoisomerase II allows the nucleosome tether to partial or fully resist supercoiling-induced DNA compaction. Figure reprinted and adapted with permission from Le et al., 2019^1^.

### Continuous winding experiments of single fibers on an MT

The magnetic tweezers data is converted into extension, turn, and force using LabView-based and MATLAB-based data conversion software. To quantify the nucleosome fiber’s quality and size, we first obtained the winding curve of the fiber under 0.5 pN tension (Figure 6C). Similar to the twisting data from angular optical tweezers, the minimal difference seen between the forward and reverse winding curves indicates the twisting process is in equilibrium without histone dissociation.

Subsequently, to study the activity of topoisomerase II on single nucleosome fiber, we introduced topoisomerase II at pM-ranged concentrations and continuously added turns up to +1000 turns with a rate of 3.6 turns/s. Figure 6D represents typical extension vs. number of turns added for a single chromatin fiber in the presence of topoisomerase II. By titrating the topoisomerase II concentration from 0 to 5 pM, it was observed that the chromatin tether can increasingly resist collapsing to the surface and the fraction of traces that can survive the entire 1000 turns increases accordingly. The characteristic topoisomerase II concentration that yields 50% full relaxation was measured to be ∼ 6 pM for single nucleosome fiber. We similarly determined that pM-ranged topoisomerase II concentration was sufficient to effectively relax naked DNA^12,13^. This observation suggests our improved surface passivation protocol for chromatin may also reduce the loss of topo activity due to non-specific surface absorption.

## H. QUANTIFICATION AND STATISTICAL ANALYSIS

The chromatin fiber tethers that pass the criteria mentioned below can be considered a tether with minimal surface interaction and nucleosome quality with known number of nucleosomes with high confidence^1,13^. These tethers can be used to perform experiments that further probe the properties of chromatin fibers and/or their interactions with other biomolecules such as topoisomerases.

### Chromatin fiber quality quantification for chromatin twisting/stretching on optical tweezers

A twisting/stretching trace is selected for a chromatin fiber with good nucleosomal content following a set of combined criteria:

1. In the stretching experiment performed subsequent to the twisting experiment, the measured contour length of DNA after nucleosome disruption at 40-45 pN must agree to a few percent of the theoretical value for that of a 12,667 bp DNA construct (Figure 6B).
2. In the stretching experiment performed subsequent to the twisting experiment, analysis must show that |*N*_in_-*N*_out_|/*N*_in_ ≤0.15, where *N*_out_ is the number of outer turns released and N_in_ is the number of inner turns released. *N*_out_ and *N*_in_ are calculated from the as described in Figure 6B.
3. In the twisting experiments, the mean difference in extension between the hat curves of adding turns and removing turns must be < 50 nm (Figure 6A).

### Chromatin fiber quality quantification for chromatin twisting on magnetic tweezers

A twisting trace is selected for a chromatin fiber following a set of combined criteria:

1. In the twisting assay performed before experiments, the mean difference in extension between the hat curves of adding turns and removing turns must be < 50 nm (Figure 6C).
2. From the hat curve’s maximum extension (extension at zero turns), we calculated the number of nucleosomes on the substrate using the linear relationship between the extension at zero turns versus number of nucleosomes established on the angular optical tweezers (Figure 7A). We selected traces with an extension consistent with 50 ± 6 nucleosomes.
3. We require the hat curve’s (+) transition width *w*_t_^+^ to be within 20% of the expected value established on the angular optical tweezers (Figure 7B). This procedure removes tethers that were partially stuck to the surface because they will exhibit short and narrow hat curves.

**Figure 7:**
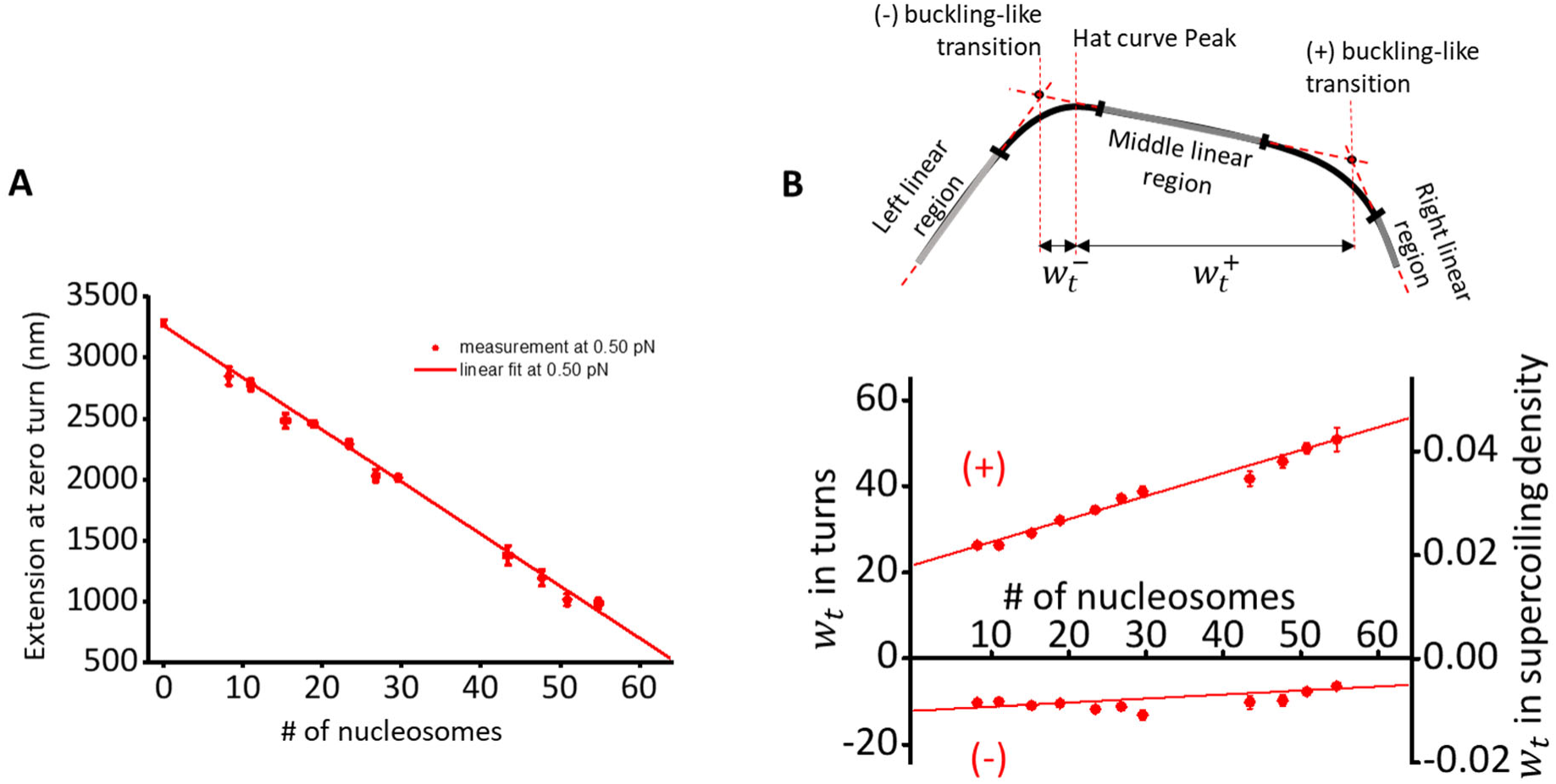
(A) Extension versus nucleosome array saturation relationships at 0.5 pN obtained from the nucleosome array stretching data on angular optical tweezers. During a twisting experiment, we use the extension at zero turns to estimate the number of nucleosomes on a substrate. The relationship at 0.5 pN is then used to estimate the number of nucleosomes on single chromatin fiber substrates in the experiments performed on magnetic tweezers. (B) Characterizing buckling-like transitions in chromatin. Each hat curve is fit by a 5-piecewise function, which consists of 3 linear regions (left, middle, and right linear, gray) and 2 quadratic regions (black). To reduce the size of the parameter space, we require that the function and its derivative be continuous. We define the position of a “buckling-like” transition to be at the intercept of the fit to the middle linear region with that of an adjacent linear region. A useful quantity to characterize these transitions is the number of turns required for the onset of the transition in either the (-) or (+) direction. Here, we call this quantity the transition width *w*_*t*_^-^ for the (-) transition, and *w*_*t*_^+^ for the (+) transition. The bottom panel shows the transition width as a function of the number of nucleosomes on DNA obtained on angular optical tweezers. Figure reprinted and adapted with permission from Le et al., 2019^1^.

## I. LIMITATI1ONS

The surface passivation method described in this manuscript has been validated in single-molecule tweezers studies of nucleosome fibers containing up to ∼ 60 nucleosomes and topoisomerase II. Application and the performance of the methods for other biomolecules awaits further investigations.

During the continuous winding experiments of nucleosome fibers on magnetic tweezers, because all tethers are constantly under a small stretching force, nucleosomes are slightly destabilized, possibly leading to a minor decrease in the number of nucleosomes within the fiber. In addition, while the mean topoisomerase II activity rate in a continuous winding experiment can be inferred from the relationship between fully relaxed fraction and the topoisomerase II concentration, the relaxation rate cannot be directly measured with this experiment^1^. For direct measurement of topoisomerase relaxation rate on a chromatin substrate, we recommend a more standard assay using magnetic tweezers where the magnetic bead does not rotate. The relaxation rate can then be obtained from the extension vs supercoiling state of a chromatin fiber^13^.

## J. TROUBLESHOOTING

### Problem 1 (related to the Section F “Forming single fiber tethers with nucleosome arrays for optical tweezers or magnetic tweezers” under STEP-BY-STEP METHOD DETAILS)

Fluid evaporation in the sample chamber preventing buffer flow.

### Potential problem

If the fluid level recedes from the sample chamber entry point due to evaporation, it can become impossible to flow in additional buffers due to the formation of a bubble that blocks fluid flow. In this case, additional fluid can be added to the exit side of the sample chamber to restore the fluid level on the entry side by the capillary effect. Furthermore, during long incubations such as the 1+ hour initial beta-casein incubation, extra fluid can be left outside of the chamber at entry/exit instead of flushing all the way through so that if some fluid evaporates, it will not affect the fluid level inside the chamber. In addition, the flow cell can be kept in a clean tip box filled with some water during long incubations to reduce evaporation.

### Problem 2 (related to the Section F “Forming single fiber tethers with nucleosome arrays for optical tweezers or magnetic tweezers” under STEP-BY-STEP METHOD DETAILS)

Cannot flow buffer into the sample chamber due to excessive hydrophobicity.

### Potential Solution

If the sample chamber becomes too hydrophobic, the capillary effect cannot pull a solution into the sample chamber. First, to avoid an excessive degree of hydrophobicity, the final sample chamber storage at step 14 under the section “Nitrocellulose flow chamber preparation” should not be more than 4 days. Alternatively, one can introduce 0.1% Tween-20 during the first flow-in step to drive the buffer into the chamber as the Tween solution has a high wettability on a hydrophobic surface^33^. Then remove the excess Tween-20 in the sample chamber by flowing in a copious amount of PBS 1X and conduct the experiments as usual. One may also construct a sample chamber with a small inlet that fits the pipette tip which can then be used to push the buffer into the chamber^34^.

### Problem 3 (related to the Section F “Forming single fiber tethers with nucleosome arrays for optical tweezers or magnetic tweezers” under STEP-BY-STEP METHOD DETAILS)

Nucleosome fiber sticking to the surface.

### Potential Solution

It is critical to use the highly purified beta-casein for surface passivation as described in this paper. Mixed casein or other casein isoforms can yield less than ideal nucleosome stretching curves, indicative of a higher degree of non-specific surface adsorption. In addition, streptavidin-coated magnetic beads and cylinders should be stored in a storage buffer with bocking proteins such as BSA and/or beta-casein.

### Problem 4 (related to the Section F “Forming single fiber tethers with nucleosome arrays for optical tweezers or magnetic tweezers” under STEP-BY-STEP METHOD DETAILS)

Nucleosome fiber losing histones after anchoring to magnetic bead.

### Potential Solution

It is critical to use the Dynabeads MyOne Streptavidin T1 magnetic bead when working with nucleosome fibers. We found that other streptavidin-coated magnetic beads, such as Dynabeads MyOne Streptavidin C1 from Thermo Fisher or S1420S from NEB, could induce histone loss after anchoring the bead to the nucleosome fiber tether. This is potentially due to an iron leakage from the bead core in a buffer containing EDTA.

### Problem 5 (related to the Section G “Twisting and stretching a nucleosome fiber with angular optical tweezers” under STEP-BY-STEP METHOD DETAILS)

Cylinder slipping on AOT when tethers are wound down close to the coverslip surface.

### Potential solution

Due to increased fluid drag on the cylinder as it approaches the coverslip surface when twisting a chromatin fiber tether, the cylinder can slip from the trap as turns are applied. To avoid this, the tether winding can be stopped and reversed once the cylinder is < 300 nm from the surface instead of applying the full -35 or +70 turns.

## K. RESOURCE AVAILABILITY

### Material Availability

Further information and requests for resources and reagents should be directed to and will be fulfilled by the Lead Contact

### Data and Code Availability

Further information and requests for data and code should be directed to and will be fulfilled by the Lead Contact

## ACKNOWLEDGEMENTS

We thank members of the Wang Laboratory for helpful discussion and comments; J.E. Baker for initial MT data analysis software; J.L. Killian for the MT data analysis software; L.D. Brennan for purification of histones; R.P. Badman for synthesizing streptavidin-coated quartz cylinders; and J. H. Lee and J.M. Berger for purification of *S. Cerevisiae* topoisomerase II. This work was supported by HHMI and NIH (T32GM008267 and R01GM136894) to M.D.W.

## AUTHOR CONTRIBUTIONS

T.T.L.: conceptualization, data curation, formal analysis, investigation, methodology, software, validation, visualization, writing – original draft; X.G.: formal analysis, investigation, methodology, software, validation, writing – review & editing; S.P.: data curation, formal analysis, investigation, methodology, validation, writing – review & editing; J.L.: data curation, validation, writing – review & editing; J.T.I.: methodology, software; M.D.W.: conceptualization, formal analysis, investigation, methodology, visualization, writing – review & edit, supervision.

